# Telomere-to-telomere assemblies reveal complex adaptive variation of 3-ketoacyl-CoA-synthases in *Populus trichocarpa* likely driven by helitrons

**DOI:** 10.1101/2025.07.10.664019

**Authors:** David Kainer, Stanton Martin, Daniel Hopp, Sophie Mosher, Timothy J. Tschaplinski, P. Doug Hyatt, Madhavi Z. Martin, Jared M. LeBoldus, Kelsey L. Søndreli, Posy E. Busby, Mengjun Shu, Kerrie Barry, Jeremy Schmutz, Anna Furches, Nan Zhao, Daniel A. Jacobson, Jin-Gui Chen, Mirko Pavicic, Priya Ranjan, Wellington Muchero, Gerald A. Tuskan, Michael R. Garvin

## Abstract

**Background:** The model woody plant *Populus trichocarpa* displays an atypical alkene-diverse wax cuticle likely driven by copy number variation (CNV) of *3-ketoacyl-CoA synthases* (*KCS*), which has been difficult to confirm based on short-read assemblies. New long-read sequencing provides opportunities to develop telomere-to-telomere resources to detect cryptic variation, including CNVs, which are currently missed in traditional analyses. Integrating this information can improve genomic prediction for breeding and provide insights into the evolutionary basis of important traits.

**Results:** Our analysis of 78 telomere-to-telomere long-read haplotypes identified more than twice as many *KCS* genes as previously reported, along with numerous intragenic non-synonymous substitutions. Random forest predictive models highlighted the importance of *Potri*.*010G079500* in producing very long chain alkenes; however, its absence did not predict previously reported alkene-deficient phenotypes. Instead, alkene levels are best predicted by the combinations of *KCS* copies. Amino acid substitutions clustered around ligand and donor binding pockets, suggesting they contribute to differing wax cuticle composition. Finally, each *KCS* gene and copy was linked to a helitron transposon. A phylogenetic analysis indicates they are the evolutionary mechanism for generating *KCS* tandem arrays.

**Conclusions:** Long-read sequencing and telomere-to-telomere assembles revealed large-effect loci critical to genetic studies that are unattainable from short-reads. These approaches also have the potential to reveal novel insights into genome structure and function, such as the helitrons identified here. Our results highlight that, given current challenges in annotation and assembly, detailed and focused long-read sequences are key to interpreting complex genomic regions that contain tandem copy number variants.

## Introduction

The leaf cuticle, which is important for managing water use and colonization by bacteria and fungi [1,2], contains an unusually large proportion (30-40%) of very long chain alkenes in *Populus trichocarpa* (Black cottonwood). Recent studies have shown that wax levels and composition change during leaf development. Interestingly, in some *Populus trichocarpa* individuals, these alkenes are completely or nearly absent across all developmental stages [3]. This developmental variability could result from 1) phenotypic plasticity due to transcriptional regulation or 2) genomic/evolutionary adaptation. A previous transcriptomic analysis of *P. trichocarpa* genotypes, displaying different developmental profiles of alkenes, identified a cluster of *3-ketoacyl-CoA synthase* (*KCS*) genes that were significantly associated with the wax content of leaves, leading to the hypothesis that natural selection has driven the adaptive alkene phenotype. This locus has also been linked to several important traits, including bud flush, growth, and disease resistance [3–5].

The KCS enzyme is a substrate-specific protein within a multi-protein complex that iteratively elongates very long-chain fatty acids (VLCFA), subsequently leading to a pathway that forms long chain alkenes [6]. Guo et al. [6], identified *Potri*.*010G079500* as a likely candidate gene for the presence and absence of alkenes in *P. trichocarpa* wax cuticle, along with four highly similar *KCS* genes (*Potri*.*010G079700, Potri*.*010G080000, Potri*.*010G080200*, and *Potri*.*010G080400*) arranged in tandem array. Notably, *P. trichocarpa* has a high redundancy of *KCS* genes compared to other species, which may have arisen from a whole-genome duplication event or tandem duplications (local copy number increase). Accurate detection and verification of copy number variation (CNV) is challenging with short-read sequencing due to read mis-mappings [7,8]. Pan-genomic approaches, which survey genomic changes across an entire species, seek to integrate allelic variation that were excluded in single reference genomes. These previously undiscovered alleles hold the potential to disentangle the evolutionary history of genes as well as generate crops with desirable traits [9–11].

As part of our efforts to clarify the KCS locus in *P. trichocarpa* PacBio sequencing was used to create telomere-to-telomere assemblies for 39 diverse genotypes (78 haplotypes). The selected genotypes vary in their cuticular alkene production, with some producing none, and others that produce intermediate and high levels. Here we report the precise map of the *KCS* gene variants and associate them with alkene variation. Our analyses revealed more than twice as many *KCS* genes, including both CNVs and allelic variants. Integrating these CNVs and new allelic variants improved our ability to predict leaf alkene content. Placement of amino acid changes on AlphaFold-generated structures indicate that the adaptations alter donor and ligand binding pocket affinities. Surprisingly, our efforts also revealed that helitron transposons may be the evolutionary mechanism driving copy number increases. Finally, while this study makes use of telomere-to-telomere assemblies, its intensive attention on a single key locus reveals the importance of long-read sequencing and the insights that can be gained by a focused, in-depth examination of a pan-genome.

## Results

### KCS genes and helitron transposons

The *P. trichocarpa* v4.1 ‘Nisqually-1’ reference genome contains seven *KCS* genes within a 350-kb region (10,400,000-10,785,000) on chromosome 10, comprising five *PotriKCS1* and two *PotriKCS2* genes. Here, using telomere-to-telomere haplotype assemblies derived from long-read sequences from 39 *P. trichocarpa* accessions, we identified 14 different *PotriKCS1* genes (henceforth referred to as *KCS*) within this locus, seven of which are copy number expansions of *Potri*.*010G080200* (Fig. 1A, Table S1a). The *KCS* sequences were classified as intragenic allelic variants if they carried variable sites but remained in the same genomic location. If the *KCS* gene differed both in sequence and genomic location, it was named a copy number expansion and assigned an alphabetic suffix. The size of *KCS* locus varied greatly across haplotypes, ranging from 169 to 293 kb, generally correlating with the total number of *KCS* genes and gene copies present, which varied between four and ten genes (Table S1b). Among the 39 genotypes, five individuals carried a nonfunctional *Potri*.*010G079500* gene, where the 5’ end of the gene had been partially copied and inserted into the opposite strand, deleting the 3’ end of the gene (Fig. 1B). These five individuals were all alkene deficient.

**Figure 1.**
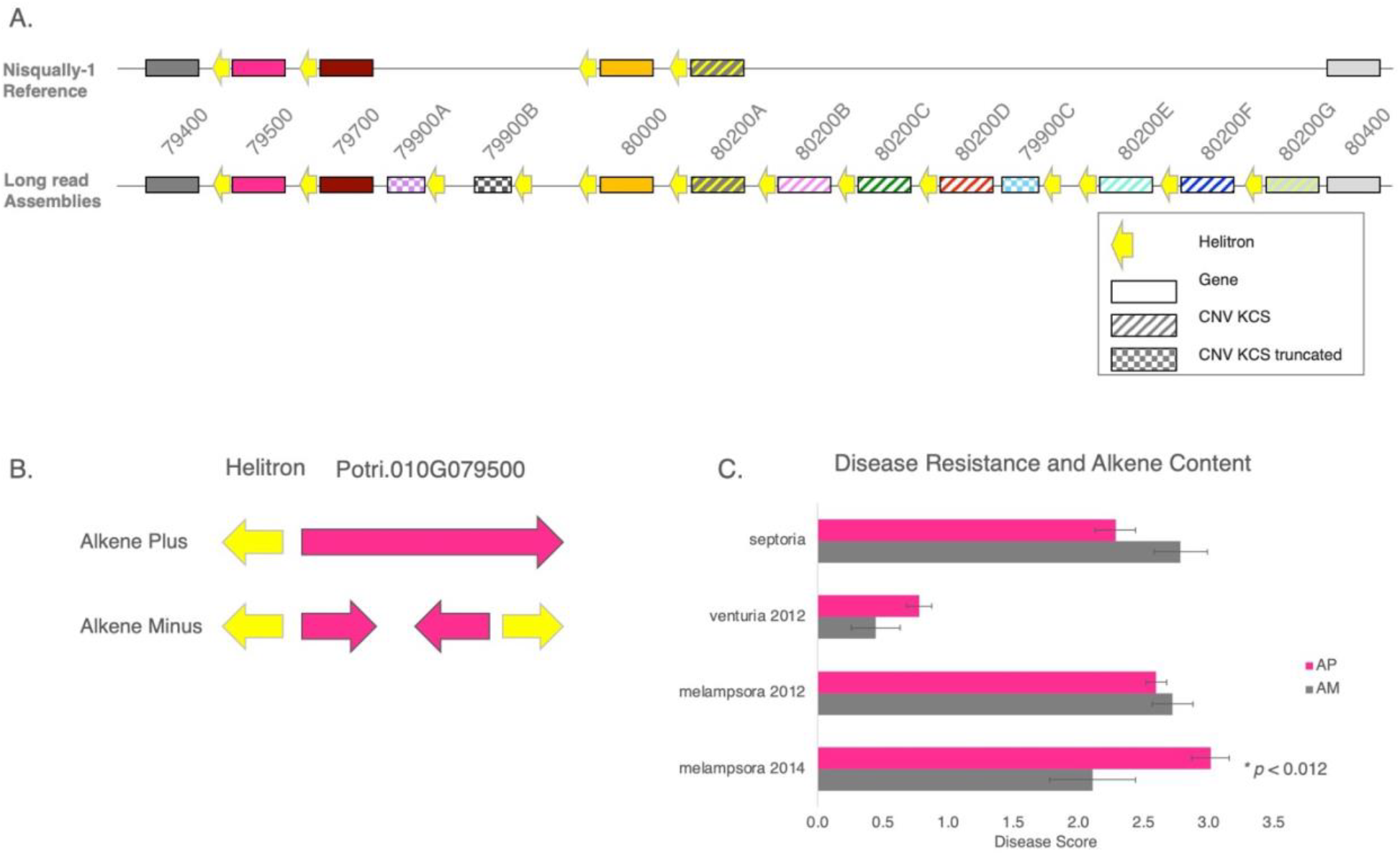
Comparison of our results to published reports of the KCS locus. **A)** KCS genes in the Nisqually-1 reference and the new 78 haploid assemblies. Solid boxes are genes (same position across genomes), and patterned boxes are copy number variants (different positions across genomes), and checkered boxes are truncated genes. Nomenclature corresponds to Nisqually-1 reference. Yellow arrows indicate helitrons with archetypal CTAG palindrome. Note, each gene is paired with a helitron. **B**) Gene model for Potri.010G079500 in alkene-minus (AM) and alkene plus (AP) haplotypes. In AM individuals, the 5’ end of Potri.010G079500 was inverted and placed on the reverse strand, deleting portions of the coding sequence, confirming the prediction of Gonzales-Vigil et al. **C**) Differences in disease severity in AM and AP phenotypes. Contrary to Gonzales-Vigil we did not find differences in S. musiva (Septoria) severity, but did find decreased severity of Melampsora 2014 (p<0.012) but not in 2012.

Previous reports identified a high number of helitrons within the *KCS* locus [3]. Our analysis revealed that each *KCS* gene and gene copy is paired with a helitron transposon, suggesting that *KCS* evolution and helitron occurrence are co-occurrent (Fig. 1A). We initially identified 11 gene-helitron pairs, but three helitrons lacked a paired gene. A subsequent search for open reading frames identified three additional *Potri*.*010G079900* copy number variants, resulting in a total of 14 *KCS* genes. The original *Potri*.*010G079900* gene, reported in the Nisqually-1 v3.0 reference, lacked a stop codon and was subsequently removed in v4.1. Additionally, the v3.0 reference indicated that *Potri*.*010G080000* contained three exons, which was reduced to two exons in v4.1. Our findings confirm that v3.0 annotation was correct, with this gene containing three exons (Table S1a). Note though that among the 39 genotypes several *KCS* genes, including those in Nisqually-1, have an indel that lacks the first exon.

To investigate potential co-evolution of helitrons and the *KCS* genes, we generated neighbor-joining (NJ) trees for the genes and their paired helitrons to determine if they were congruent (Fig. 2). The NJ trees, based on genes, and the tree based on the paired helitrons, were highly congruent with regard to topology, clustering by *KCS* gene, and copy number type. However, in two cases, a helitron from one gene was paired with a different *KCS* gene. For example, the helitron paired with the *Potri*.*010G079500* gene in haplotype BESC-377_H2 was nearly identical to the helitron preceding *Potri*.*010G079700* in that same individual. Additionally, the upstream sequence contains several other unique features, including a copy of a chloroplast-derived *DRT111* gene, linked to adaptive traits from a previous genome-wide association study [4]. This unique upstream sequence also contains two transposases, one of which harbors an “HUH” domain that is indicative of helitron transposition machinery [12,13]. This transposase is linked to the *Potri010G079700* gene across all haplotypes, *i*.*e*., haplotypes lacking the *Potri010G079700* gene are either missing this transposase or have it partially deleted (Table S2). Finally, contrary to previous reports, our results indicate that these alkene deficient genotypes are not more susceptible to *Sphaerulina musiva* compared to the full-length wildtype. They appear resistant to the leaf pathogen *Melampsora*, although our sample size is relatively small (Fig. 1C, Table S3).

**Figure 2.**
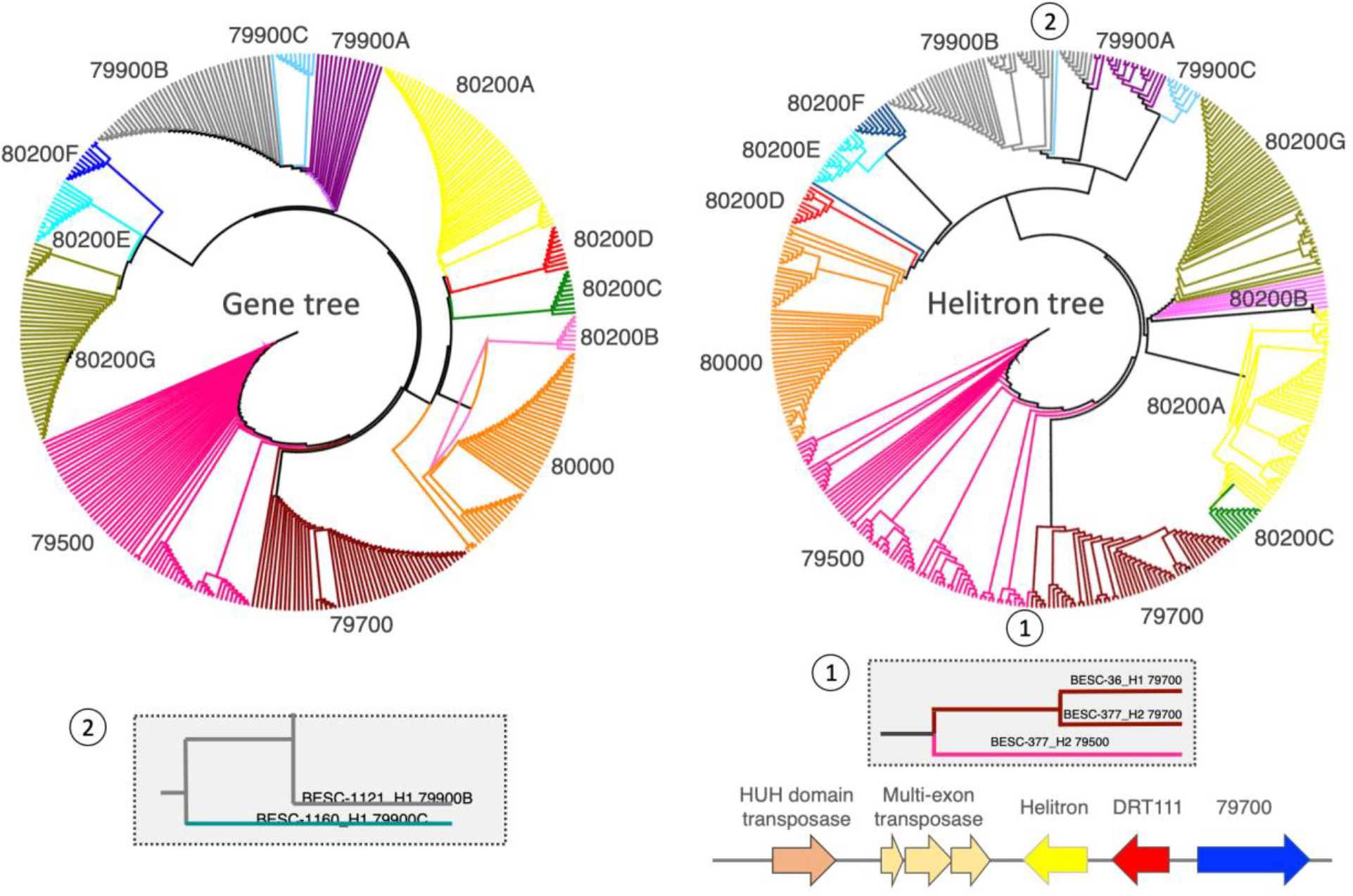
Neighbor-joining (NJ) trees based on exons of the genes and copy number variants (left) compared to NJ tree based on their paired helitrons (right). Colors correspond to those in Figure 1A. Helitron sequences are mostly conserved, and the topology is congruent with the exon-based tree, but there are two major discrepancies (noted with numerals 1 and 2). The coding sequences of Potri.010G079500 and Potri.010G079700 in ‘BESC-377’ vary, even though their helitrons are nearly identical. Likewise, the helitron sequence from BESC-1160 paired with Potri.010G079900C is most similar to that of BESC-1121 for Potri.010G079900B even though the gene sequences are different. The sequences shared between Potri.010G079500 and Potri.010G079700 are unique in that they both harbor a chloroplast-derived gene (DRT111) and two transposases. This DRT111-transposase sequence is found in all haplotypes with a full-length Potri.010G079700 gene. In BESC-377_H2 it also appears upstream from Potri.010G079500, suggesting a shared evolutionary history.

### Predictive models for alkene abundance using the KCS locus

First, we predicted the binary trait of alkene-minus (AM) or alkene plus (AP) phenotypes by fitting an iterative Random Forest (iRF) classification model to determine which gene variants displayed the greatest importance. *Potri*.*010G080200* copy number variants *80200B* and *80200G_A58V* emerged as the most important predictors (Fig. 3, left). The Shapley analysis revealed that any non-zero dosage of *Potri*.*010G079400* or *Potri*.*010G079500* strongly predisposes individuals towards the AM class. Contrary to expectations [3], *Potri*.*010G079500* did not show significant predictive importance. To clarify this observation, we fit iRF models to explain the variation in abundance of Z-9-pentacosene and Z-9-heptacosene, focusing only on individuals that produce the alkenes (i.e., AM individuals were excluded from this analysis). A wide variety of *KCS* genes and variants were found to be important in these models, including genes of both *KCS1* and *KCS2* classes (Fig. 3, center and left). However, only some genes were consistently important across both alkenes. That is, multiple variants of the gene *Potri*.*010G079500* were important for one alkene or the other, but are clearly not the only drivers of alkene abundance variation at this locus. For example, having a double variant, 79400_E208D_C283R, in the *Potri*.*010G079400* gene, a more distantly related class 2 *KCS* gene [3], corresponds to lower levels of both hepta- and pentacosene.

**Figure 3.**
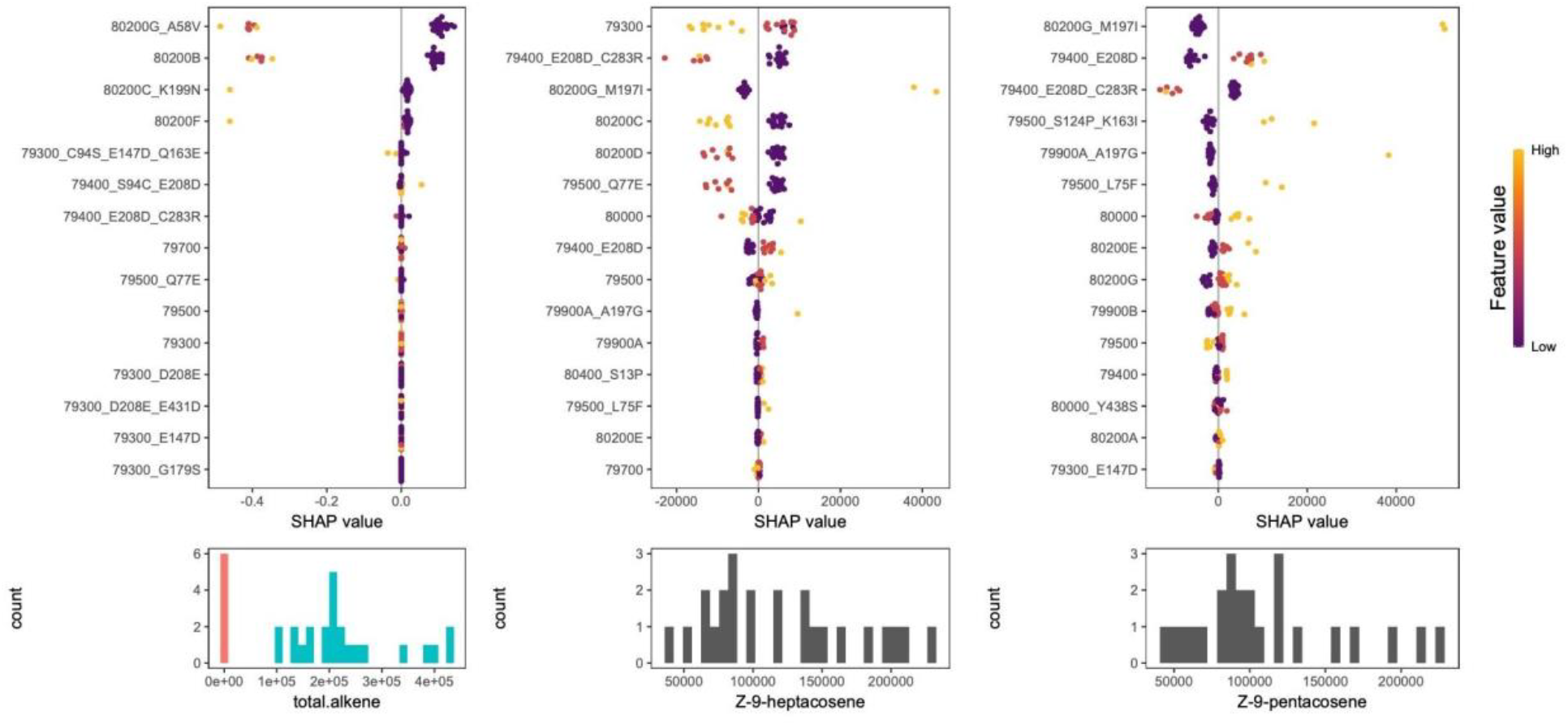
Explainable AI analysis of iterative Random Forest (iRF) models for different alkene phenotypes. The y-axis represents the SHAP value of each individual gene or copy number variant. Each filled circle represents an individual and the color represents the feature values, which is the number (dosage) of that feature for that individual. Purple indicates 0 alleles and yellow is maximum number of alleles for a gene variant (usually two, but can be one). Histograms below each SHAP plot depict the phenotype’s distribution and SHAP values represent the tendency for that feature-allele combination to drive the phenotype prediction for an individual towards either low (left side) or high (right side) values. The model to predict alkene deficient (AM, red bars) versus alkene plus (AP, blue bars) phenotypes indicate 80200B and 80200G_A58V are the most accurate predictors for predicting AM or AP class (left plot), where high dosage (1 or 2) strongly drives individuals towards the AM class. The models that predict heptacosene (center) and pentacosene (right) identify a variant of the type 2 KCS protein Potri.010G079400 as important (E208D) for both but addition of the C283R mutation decreases these alkenes. The presence of Potri.010G080000 has opposing effects on the two alkene types and different variants of the Potri.010G079500 are important for predicting pentacosene levels but not heptacosene.

### Structural analysis

The first 106 amino acids of the AlphaFold prediction are disordered, making it unclear how amino acid changes here affect function. Supporting the hypothesis of adaptive selection of *KCS* genes, more than half (16/25) of the amino acid changes outside the disordered region cluster around the ligand binding pocket, with a few variants near the donor binding site for malonyl-CoA (Fig. 4, Table S1). Many of the substitutions are predicted to alter hydrogen bonds, potentially impacting the active site directly. For example, the intragenic variant Y438S in Potri.010G080000 is predicted to alter a hydrogen bond with E433, re-positioning the catalytic residue 444N (site 456 in Figure 4) at the other end of the alpha helix on which it sits.

**Figure 4.**
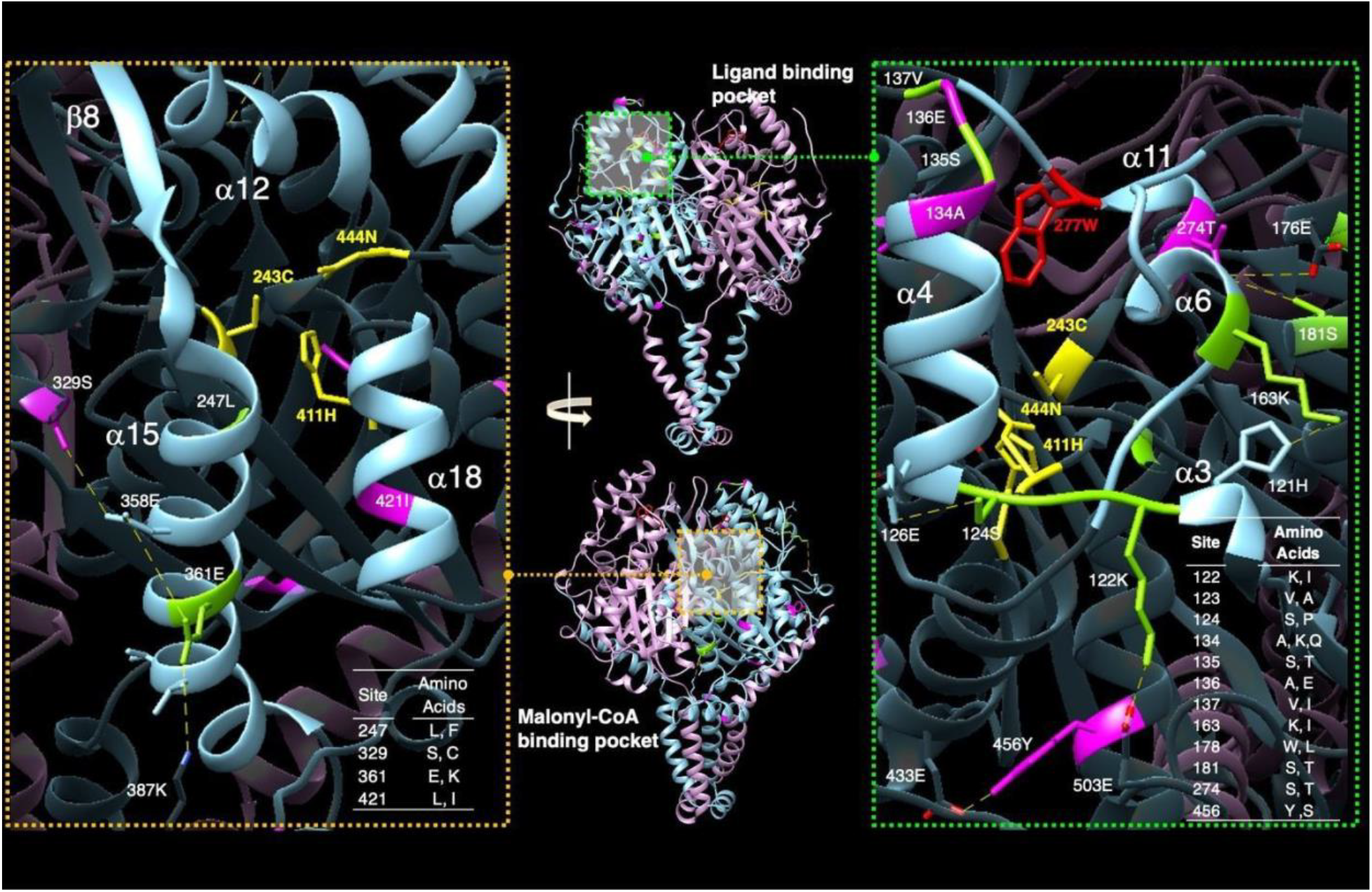
Amino acid substitutions overlaid on the AlphaFold predicted structure for Potri.010G080200. Previous work identified the binding pockets for the ligand (top, center panel) and the two-carbon donor malonyl-CoA (bottom, center panel). Substitutions among KCS variants in general (green residues in left and right panel) and those specific to the Potri.010G080200 copies (pink residues left and right panels), cluster around the ligand binding pocket (magnified view with light blue coloring, right panel). The tryptophan at position 277 (as shown in red in the right panel) is necessary for function, active Cys-His-Asn triad (in yellow). Substitutions are also found in the malonyl-CoA binding pocket (magnified in the left panel). Changes are predicted to alter the hydrophobicity and shape of the region and likely affect ligand binding/activity (malonyl-CoA binding).

## Discussion

Our comprehensive long-read, telomere-to-telomere analysis emphasizes the importance of obtaining accurate genomic information in linking genes to phenotypes. The results we present here partially support and partially refute previous reports on the *KCS* locus as it relates to very long chain alkene content of the leaf cuticle. On one hand, we provide a mechanistic explanation for the observation that as a group, individuals that produce very long chain alkenes (referred to as AP) have two functional copies of *Potri*.*010G079500*. In agreement with Gonzales-Vigil *et al*. [3], five of our six alkene deficient individuals (referred to as AM) have a structural variant in at least one haplotype, which likely results in no transcript and explains the lower mRNA transcript levels observed in their AM group as a whole. However, at the individual level, the loss of *Potri*.*010G079500* does not fully explain the AM phenotype, as four of our six AM individuals carry at least one functional copy of *Potri*.*010G079500*, and one AM individual carries two copies. A closer examination of Gonzalez-Vigil *et al*.’s data supports our observation, as two of the individuals in their AM population show expression of *Potri*.*010G079500*. However, direct comparisons are challenging since they mapped their reads to *P. trichocarpa* v3.0, lacking the detailed information we present here. In contrast, our study shows that the presence of the *KCS* copies *Potri*.*010G080200B* and *Potri*.*010G080200G_A58V* are accurate markers for genomic prediction of the AM phenotype.

An unexpected and remarkable finding was the association of a helitron transposon with each of the *KCS* genes, suggesting that helitrons play a mechanistic role in generating copies of these genes (Fig. 5). Helitrons are mobile elements with a peel-and-paste mechanism of transposition, distinct from the more commonly known cut-and-paste or copy-and-paste mechanisms of other transposons, and helitrons do not generate duplicated regions at either the 5’ or 3’ ends [14,15]. This feature makes them theoretically ideal for generating new functional copies of genes with little risk of inducing mutations. One reason they may have been missed in previous studies is that they are difficult to predict, given the lack of sequence conservation outside a 5’ CTAG palindrome and a 3’ hairpin loop at the opposite end. In addition, the multiple copies of the helitrons across all loci complicate read mapping using short-read sequences. Given all the above, our hypothesis that helitrons are responsible for generating copies of *KCS* and which is supported by reports of helitron-mediated gene transfer in maize [16,17]

**Figure 5.**
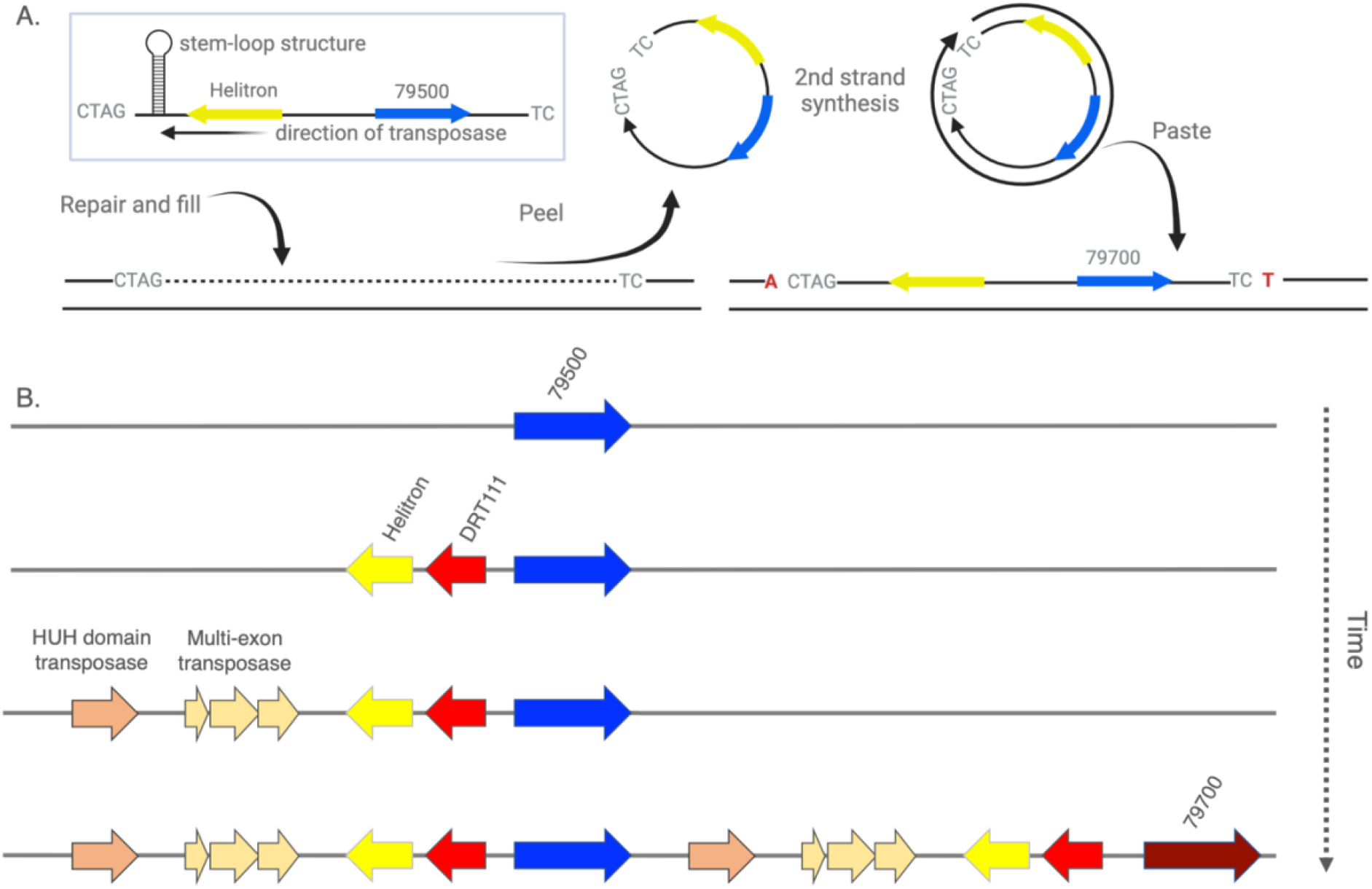
**A)** Helitrons have a unique peel-and-paste method of transposition in which a rolling circle is generated and inserted between an AT target nucleotide pair, with no duplications at the end, and the empty single strand from which it was derived is restored, retaining the original gene. Nearly all helitrons harbor an ‘A’ upstream from the ‘CTAG’ site, and a ‘TCT’ just downstream from its KCS paired gene (Table S6, S7), as would be expected if they were inserted as helitrons. Several alkene-deficient genotypes have mutations in the stem-loop structure that may affect helitron efficiency. B) Hypothetical evolutionary model of copy number generation via helitrons. Given the shared helitron between Potri.010G79500 and Potri.010G079700 in BESC-377_H2 and the presence of that helitron in all Potri.010G79700 sequences, we propose that Potri.010G079500 represents the origin of KCS copies. Once the transposase machinery and the DRT111 gene were inserted proximal to the helitron in Potri.010G79500 it was able to replicate itself and generate Potri.010G79700. This additional gene copy provided an advantage and rapidly spread throughout the population.

The majority of helitrons reported in the literature are non-autonomous and require transposase machinery for their movement. In plants, these enzymes typically feature an “HUH” domain and an associated helicase domain, often with single stranded repair activity. It is tempting to speculate that the transposases and the DRT111chloroplast derived DNA repair genes linked to the *Potri*.*010G79700* gene could be the “helicase” machinery in question in *P. trichocarpa*. Regrettably, the AlphaFold database currently lacks predicted structures from these mobile elements, making it difficult to computationally determine if any of these transposases could perform that function.

## Limitations of our study

The results we presented here required a considerable effort for manual curation and annotation. Although algorithmic approaches can easily identify presence/absence and copy number counts for downstream analyses, to our knowledge, the only means to extract intragenic allelic variation and features such as helitrons is through manual exploration of each individual haplotype. For instance, changes in splice sites generated an incorrect annotation of *Potri*.*010G80000* and removal of the *Potri*.*010G79900* gene in v4.1 of the *P. trichocarpa* genome. Our gene models, while predicted, will need to verification with transcriptome data, preferably also using long-read sequencing. Likewise, experimental evidence will be necessary to unravel the complexities arising from the combinatorial contribution of all KCS proteins within an individual. In *Arabidopsis*, it has been shown that KCS proteins act as dimers that result in different products [18,19]. If the same is true for *P. trichocarpa*, this would indicate 105 possible hetero- and homodimers that may produce a highly varied and complex combination of very long-chain fatty acids.

Another limitation of this study was our small sample size (N=39 individuals with 30 having metabolomic data). Still, with the depth of information that we were able to extract from these individuals, we added considerable knowledge to the genetic basis and evolutionary underpinnings of alkene production in *P. trichocarpa*. This new information indicates that production of the wax cuticle is much more complicated and will require a deeper examination of the underlying biology. Our work leads to several questions; are all of the *KCS* alleles in an individual expressed and translated to protein or is there allele-specific expression? How do these different variants interact with each other and other proteins in the fatty acid synthesis pathway and what is the full spectrum of fatty acids that result from those interactions? Is the effect on different phenotypic traits such as growth and disease resistance directly related to the wax cuticle composition or is that simply a correlated phenotype, and is the ultimate effect due to upstream changes in fatty acid compositions such as oleic acid? These questions cannot be addressed with short read shotgun approaches, but rather will require long read RNAseq and proteomics. Lastly, systems will need to be developed for naming these new gene variants. Intragenic variation is easily handled as we have done here, but copy number represents a new challenge and will likely require additional verification efforts and new naming rules.

## Methods

### Source material of P. trichocarpa

Plant material from 1,081 *Populus trichocarpa* genotypes, originally collected from wild populations in California, Oregon, Washington, and British Columbia were planted in a stool bed at the Oregon State University Research Farm in Corvallis, OR [4]. During January 2014, dormant branch cuttings were collected and sent to the North Dakota State University’s Agricultural experiment station research greenhouse complex in Fargo, ND. For each genotype, branches were cut into 10 cuttings, measuring 10 cm in length, with at least one bud. Cuttings were soaked in distilled water for 48 h, planted in conetainers (Ray Leach SC10 Super Cone-tainers, Stuewe and Sons, Inc., Tangent, Oregon, USA) measuring 3.8-cm in diameter and 21-cm deep filled with growing medium (SunGro Professional Mix #8; SunGro Horticulture Ltd., Agawam, MA) amended with 12 g of Nutricote slow release fertilizer (15-9-12) (N-P-K) (7.0% NH_3_-N, 8.0% NO_3_-N, 9.0% P_2_O_5_, 12.0% K_2_O, 1.0% Mg, 2.3% S, 0.02% B, 0.05% Cu, 0.45% Fe, 0.23% chelated Fe, 0.06% Mn, 0.02% Mo, 0.05% Zn; Scotts Osmocote Plus; Scotts Company Ltd., Marysville, OH). The cuttings were planted such that the upper most bud remained above the surface of the growing medium. Plants were grown in a greenhouse with a temperature regime of 20°C/16°C (day/night) and an 18-h photoperiod supplemented with 600 W high-pressure sodium lamps. Slow-release fertilizer was added weekly with 15-30-15 (N-P-K) Jack’s fertilizer (Jr. Peters Inc; Allentown, PA) at 200 ppm for two months to promote root growth and subsequently fertilized with 20-20-20 (N-P-K) liquid fertilizer (Scotts Peters Professional; Scotts Company Ltd., Marysville, OH) once a week. Plants were watered as needed.

### Sequence analysis

We extracted the 250-kb region that included the KCS locus from 39 chromosome resolved, haplotype phased, PacBio HiFi integrated genome assemblies provided by the Joint Genome Institute and Hudson Alpha (39 genotypes, 78 haplotypes). The first 30 nucleotides beginning from the ATG start codon are conserved in all five KCS genes (*Potri*.*010G079500, Potri*.*010G079700, Potri*.*010G080000, Potri*.*010G080200, Potri*.*010G080400*), and therefore, this sequence string was used to search for all possible *KCS* genes in each haplotype in the 250-kb region of chromosome 10. We then extracted 1812 base pairs (the longest gene of the five), produced a multiple sequence alignment, removed the intron based on the intron/exon structure defined by the Nisqually-1 reference, and translated the sequences. Pseudogenes were annotated as “Nonsense” versions of the gene, which included frame-shift mutations and large deletions. Once each haplotype was annotated, we scored each for presence/absence of all known *KCS* genes (0/1). We then combined haplotypes to generate diplotypes and summed the two scores to generate a 0/1/2 genotype. Neighbor-joining trees were generated in CLC Genomics Workbench and visualized with FigTree v1.4.4. Branch lengths represent bootstrap values. Helitron transposons were identified with a BLAST search of the sequences taken from the repeatmask track of Nisqually-1 v4.1 against each haplotype.

### Metabolite phenotypes

Fully expanded leaves (leaf plastochron index 9±1) were collected for 851 black cottonwood poplar (*P. trichocarpa*) genotypes in genome-wide association studies (GWAS) at Clatskanie, OR over three consecutive sunny days in July 2012, as described previously [20]. Briefly, aqueous ethanolic extracts (80%) were analyzed for metabolites by gas chromatography-mass spectrometry following trimethylsilylation. Both automated and manual data extraction were used to extract metabolite peaks, which were then normalized to the amount of internal standard (sorbitol) injected and mass of leaf extracted.

### Disease phenotypes

Disease severity, caused by *Melampsora* spp. and *Venturia populina* in *P. trichocarpa*, was evaluated at the research plantation in Clatskanie, OR. *Melampsora* spp. infections were measured in 2012 and 2014 using a scale where 0 indicates no rust signs, 1-very light rust occurrence, 2-rust on most leaves in some small clusters, 3-rust dominant on most leaves, but no leaf necrosis, and 4-heavy rusting with necrotic spots on leaves. For *Venturia populina*, disease severity was assessed in 2013 with a scoring system: 0 - no apparent *Venturia*, 1 - a few leaves infected, 2 most leaves infected and a small number of petioles and terminal branch buds affected, and 3 – majority of the petioles and terminal buds infected, a common shepherd’s crook appearance.

Three isolates of *Sphaerulina musiva* (MN-12, MN-14, MN-20), collected from three separate trees in near Garfield, MN, USA, were chosen for inoculation, based on preliminary virulence testing, and transferred from storage (−80^°^C) onto K-V8 (180 ml of V8 juice [Campbell Soup Company, Camden, NJ], 2 g of calcium carbonate, 20 g of agar, and 820 ml of deionized water) growth media, sealed with Parafilm (Structure Probe Inc., West Chester, PA). Petri plates were placed on a light bench under full-spectrum fluorescent bulbs (Sylvania; Osram Gmbh, Munich) at room temperature until sporulation was observed. Following sporulation, five 5-mm plugs were transferred onto another K-V8 plate and grown for 14 days under continuous light. There were a total of 200 plates for each isolate.

The experimental design was a randomized complete block design with 4 blocks. Plants were inoculated when they reached a minimum height of 30 cm (∼54 days after planting). Plates containing isolates were unsealed and ∼1 ml of deionized water was added to the plate. Rubbing the media surface with an inoculation loop dislodged the spores and the spore suspension was collected with a pipette. The spore suspensions were individually bulked from the three isolates at a concentration of 10^6^ spores ml^-1^ for each isolate. Plants were taken out of the greenhouse and their heights were measured prior to inoculation, sprayed with a HVLP gravity fed air spray gun (Central Pneumatic, Harbor Freight Tools) at 20 psi until the entire leaf and stem surface was wet (15 ml) and placed into a black plastic bag for 48 h. At three weeks post-inoculation phenotypic responses were characterized by evaluating disease severity using a scale from 1-5. (1= no disease/cankers, 2= small necrotic lesions but resistant reaction taking place, 3= small (larger than 2) with necrosis extending beyond resistant response, 4= large necrotic lesions 5= stem girdling lesions occasionally with sporulation. Tests for differences of disease phenotypes was performed with a student’s t-test.

### Genotype-to-Alkene models

Since VLC alkene production in this population appears to have a binary tendency (AM / AP), we fit an iterative random forest (iRF) to classify individuals as AM or AP based on their KCS genotypes. This model was fitted using accessions with both genotype and metabolite data using the iRF v3.0.0 package in R with 1000 trees and 5 iterations [24]. KCS genotypes were encoded as 0/1/2 for the iRF predictor matrix [22,23]. Furthermore, the AP individuals show a wide distribution of alkene abundances, so to determine which KCS genes were most associated with alkene variation we then fit an iRF model for each of Z-9-pentacosene and Z-9-heptacosene abundances using the KCS genotypes for only the AP individuals. All iRF models were performed using the iRF v3.0.0 package in R with 1000 trees and 5 iterations [24]. Importance scores for each model were then extracted from the iteration with the best fit determined by out-of-bag accuracy.

### Shapley analysis

Shapley analysis was applied to the outputs of each iRF model using the fastshap v0.1.1 package in R to explain the influence of the most important predictors on the dependent variable. Each shapley analysis used 500 Monte-Carlo repetitions. Visualizations were generated with the shapviz v0.9.3 R package.[25]

### Structural analyses

We used the AlphaFold structure U5G1R7 for *Potri*.*010G080200* (POPTR_010G080200 on AlphaFold). We aligned with the 2ix4 PDB file for *Arabidopsis thaliana* mitochondrial beta-ketoacyl ACP synthase hexanoic acid complex using the MatchMaker function in Chimera with a Smith-Waterman algorithm. Substrate and malonyl-CoA binding pockets as well as numbering for alpha-helices and beta-sheets were taken from Chen et al.^5^. Structure is shown as a dimer as in *Arabidopsis thaliana*. Predictions were performed in ChimeraX using AlphaFold [26]

## Acknowledgements and Funding

This material is based upon work supported by the Center for Bioenergy Innovation (CBI), U.S. Department of Energy, Office of Science, Biological and Environmental Research Program under Award Number ERKP886. The work (proposal: 10.46936/10.25585/60001339) conducted by the U.S. Department of Energy Joint Genome Institute (https://ror.org/04xm1d337), a DOE Office of Science User Facility, is supported by the Office of Science of the U.S. Department of Energy operated under Contract No. DE-AC02-05CH11231. We thank the Department of Energy Joint Genome Institute and collaborators at the Center for Bioenergy Innovation for pre-publication access and use of the *KCS* gene from the ongoing poplar pan-genome project. This research used resources of the Oak Ridge Leadership Computing Facility at the Oak Ridge National Laboratory, which is supported by the Office of Science of the U.S. Department of Energy under Contract No. DE-AC05-00OR22725. This work was partly funded by The Australian Research Council Centre of Excellence for Plant Success in Nature and Agriculture (CE200100015). This work was also funded by CAREER awards NSF 2146552 and USDA 2022-67013-37437.

## Data Availability

Data for this manuscript has been released at: http//10.25983/2322563 or available in the Supplementary Tables

## Author contribution

DK – Conceptualization, Methodology, Validation, Formal analysis, Investigation, Writing Original Draft, Writing Review & Editing, Visualization; SM – Software, Data Curation, Visualization, Conceptualization, Methodology, Supervision, Writing Review and Editing; DH – Software; SM – Software, Data curation, Visualization; TT – Data Curation, Investigation; PH – Data Curation, Investigation; MM – Data Curation Investigation; JL – Data Curation, Investigation; KS – Data Curation, Investigation; PB – Resources, Writing - Review & Editing, Funding acquisition; MS – Writing - Review & Editing; KB – Investigation; JS – Investigation; AF – Data Curation, Investigation; NZ – Data Curation, Investigation; DJ – Data Curation; JC – Investigation, Resources, Writing - Review & Editing, MP – Data Curation Investigation; PR – Data Curation; WM – Data Curation; GT – Conceptualization, Data Curation, Funding acquisition, Project administration, Supervision, Project administration, Writing Original Draft, Writing Review & Editing; MG – Conceptualization, Methodology, Validation, Formal analysis, Investigation, Writing Original Draft, Writing Review & Editing, Visualization, Supervision.

